# Computational modeling highlights disordered Formin Homology 1 domain’s role in profilin-actin transfer

**DOI:** 10.1101/263566

**Authors:** Brandon G. Horan, Gül H. Zerze, Young C. Kim, Dimitrios Vavylonis, Jeetain Mittal

## Abstract

Formins accelerate actin polymerization, assumed to occur through flexible FH1 domain mediated transfer of profilin-actin to the barbed end. To study FH1 properties and address sequence effects including varying length/distribution of profilin-binding proline-rich motifs, we performed all-atom simulations of mouse mDia1, mDia2; budding yeast Bni1, Bnr1; fission yeast Cdc12, For3, and Fus1 FH1s. We find FH1 has flexible regions between high propensity polyproline helix regions. A coarse-grained model retaining sequence-specificity, assuming rigid polyproline segments, describes their size. Multiple profilins and profilin-actin complexes can simultaneously bind, expanding mDia1-FH1, which may be important in cells. Simulations of the barbed end bound to Bni1-FH1-FH2 dimer show the leading FH1 can better transfer profilin or profilin-actin, having decreasing probability with increasing distance from FH2.

## 1 INTRODUCTION

Formins are dimer-forming actin regulators that play fundamental role in biological processes such as cytokinesis, cell motility and muscle development [21, 42]. They function by nucleating new actin filaments and remaining processively associated with the filament’s barbed end during elongation. The Formin Homology 2 (FH2) domain dimerizes and wraps around the barbed end bringing with it adjacent Formin Homology 1 (FH1) domain [21, 42]. One key feature of the FH1 domain is that it is proline-rich, often containing many proline-rich motifs (PRMs). These PRMs can bind profilin, which is an actin regulator, and help accelerate filament elongation [30, 45, 52]. The length and distribution of these PRMs varies widely among formins (1-13 consecutive prolines, 2-14 potential proline-rich profilin-binding sites, 16-50% proline content for formins in this study), playing an important role in actin regulation [41, 56].

Actin filament ends associated with FH1-FH2 dimers polymerize actin at a reduced rate compared to free filaments by the “gating” factor, which varies between near zero and 1, depending on the formin [30, 52]. In the presence of profilin, polymerization is further accelerated by multiple fold [30, 45]. Kinetic modeling has suggested a “transfer” mechanism (originally proposed for actoclampin [17, 18]) by which a flexible FH1 captures profilin-actin complexes and then directly delivers them to the FH2-bound barbed end with sufficiently high probability that can overcome even small gating factors [52]. In this model, PRMs closer to the FH2 domain transfer actin at greater efficiency compared to more distant PRMs, a result supported by experiments with formins of varying FH1 length [13, 41].

Several questions remain about biophysical properties and basic function of FH1 domains. Based on the sequence composition, FH1 is expected to be disordered [24], but there is, in general, a lack of molecular characterization of its equilibrium structural ensemble and how it may be modulated by the differences in PRMs among different formins. It has been hypothesized that each FH1 of the formin dimer is specific to one of the two actin protofilaments, but how each FH1 can geometrically and physically achieve direct transfer of profilin-actin has not been studied in detail. It is also not clear how binding of one or multiple profilin or profilin-actin complexes can influence FH1 structure. This lack in understanding of basic biophysical properties of formins poses barriers to resolving many issues pertaining to their role in actin polymerization, such as the observed dependence of external force on formin-mediated polymerization [14, 27, 31, 51, 53, 56] and acceleration of polymerization in the presence of cofactors [22].

Prior computational models of the FH1 domain have been largely devoid of atomistic-level details and, therefore, cannot help with the issues raised above [11, 52]. A recent study by Zhao *et al.* [55] used an atomistic model, which uses probabilistic sampling of protein conformations as opposed to physics-based force-fields [36], to study the FH1 sequence of mouse formin mDia1. They proposed that profilin binding to FH1 causes a cooperative coil-to-elongation transition [55].

All-atom molecular dynamics (MD) simulations based on accurate physics-based force fields coupled with enhanced sampling techniques such as parallel tempering (PT), well-tempered metadynamics can provide experimentally validated structural ensembles of disordered proteins [10, 54]. Detailed structural information from these simulations can further be used to develop computationally efficient coarse-grained models to study large-scale phenomena. A recent study used all-atom MD simulations in conjunction with a coarse-grained model to study the electrostatic interactions between the FH2 domain of budding yeast formin Bni1 and the barbed end of the actin filament [3].

In this work, we examine several FH1 domains using all-atom explicit solvent simulations and show that these proteins behave as typical intrinsically disordered proteins (IDPs) and occupy high-propensity poly-L-proline (PP) helices within the PRMs. Additionally, we develop a C_α_-based coarse-grained model which retains the disordered characteristics of the FH1 [19, 35] outside the PRMs and PP within the PRMs. We use this model to study the effect of profilin or profilin-actin binding on the FH1 domain of mDia1 that has high PRM density. We find that all PRMs can be occupied and quantify the resulting FH1 expansion. To model the transfer mechanism of FH1-bound actin, we also simulate FH1-FH2 in complex with a model actin filament, using the prior model of Bni1 FH2 associated with the barbed end [3]. The computed closure rates from these simulations as a function of distance away from the barbed end agree well with previous predictions from a simple polymer theory [52]. We also find that the corresponding PRMs of the two FH1s of the FH1-FH2 dimer have different abilities to contribute to polymerization, with those close to the FH2 and nearby the polymerization site having the higher rate.

## 2 MATERIALS AND METHODS

### All-atom Explicit Solvent Model and Simulation Details

Peptides are modelled using Amber99SBws protein force field [7] and solvated in TIP4P/2005 water model [1] with 10% strengthened protein-water interactions to correct overly collapsed nature of unfolded proteins [7]. Amber99SBws force field is based on Amber99SB*-ILDNQ [8] which has a backbone correction for helix propensity [6], modified torsion parameters for some of the sidechains [33] and a unified backbone charges [8]. The mDia1 and Cdc12 peptides are solvated in a truncated octahedron box with 9.3 nm (81111 atoms) and 9.8 nm (95199 atoms) spaced faces, respectively. Initial coordinates are briefly energy minimized for 500 steps and equilibrated for 100 ps in NVT ensemble followed by 100 ps in NPT ensemble, where pressure is maintained at 1 bar using isotropic Berendsen pressure coupling [4]. Further production runs are performed in NVT ensembles. Systems are propagated using stochastic Langevin dynamics with a friction coefficient of 1/ps. Electrostatic interactions are calculated using the particle-mesh Ewald method [20] with a real space cutoff distance of 0.9 nm. A 1.2 nm cutoff distance is used for the van der Waals interactions.

Standard molecular dynamics simulations are performed at 300K for 7 FH1 domains (see Fig. 1) using GROMACS-4.6.7 [5, 23] for at least 500 ns. Simulations of FH1 domains mDia1 and Cdc12 are also done using parallel-tempering (PT) [49] in the well-tempered ensemble (WTE) technique to efficiently sample the equilibrium ensembles of these peptides. In a PTWTE simulation multiple replicas of the system are propagated in parallel at different temperatures like a standard PT simulation, but the potential energy fluctuations are amplified in the WTE thereby allowing one to use fewer number of replicas [10, 16]. We used 20 replicas which are distributed between 300 K and 517 K to obtain uniform acceptance probability of approximately 20% between all adjacent replica pairs [44]. Potential energy of the system is biased using a bias factor of 40, and a Gaussian width and height of 500 and 1.5 kJ/mol (initial) applied at every 2000 steps. PTWTE simulations are performed using PLUMED-2.1.1 [9] plugin for 220 (mDia1) and 200 (Cdc12) ns per replica. For the first 50 ns of the run, WTE ensemble is turned on for all the replicas but 300 K replica [40, 50]. Energy bias accumulated during the first 50 ns is used as a static bias for further continuation of these simulations. Analysis of the last 150 ns (300 K replica) is presented in the Results section.

**FIGURE 1.**
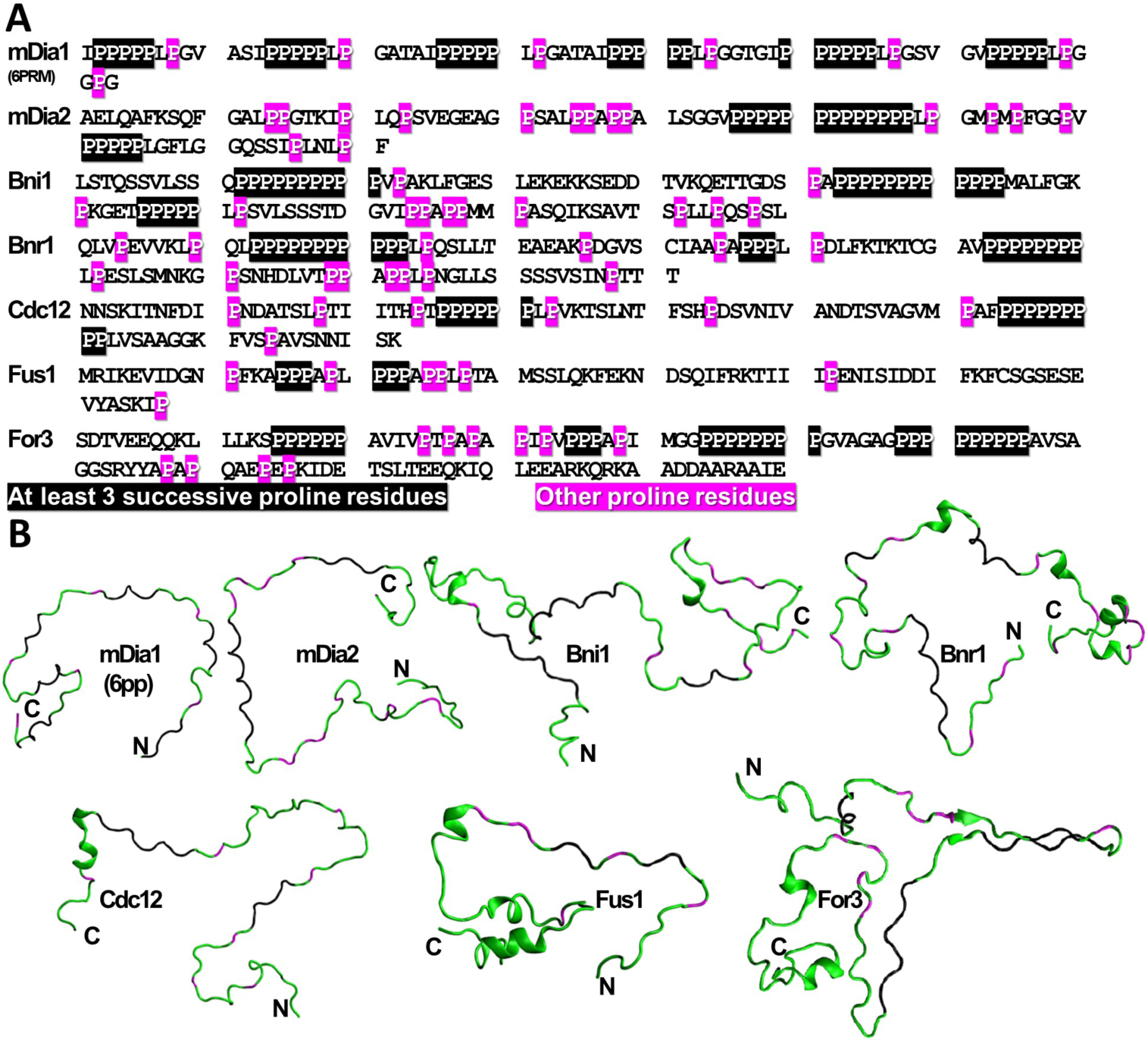
Primary structure of FH1s studied. (A) Primary structure of the FH1s studied in this work. Residues highlighted in black contain at least three successive proline residues (the same criterion was used to make PRMs rigid in the coarse-grained simulations). Residues highlighted in magenta are other proline residues. (B) Screenshots from the all-atom simulations. Proline residues colored as in (A). Other residues colored in green. Location of N- and C-termini are labeled with “N” and “C”, respectively.

### Coarse-grained Model and Simulation Details

To develop a coarse-grained description of FH1 and other proteins involved in the actin polymerization, we use the Kim-Hummer (KH) model [28] which was proposed to study weak protein-protein interactions among rigid folded proteins. Specifically, in this model, virtual bonds between neighboring C_α_ atoms of a flexible chain are represented as a harmonic potential with equilibrium distance 0.381nm. Nonbonded pairwise interactions are modeled by a standard Lennard-Jones-type potential function for all residue pairs and by Debye-Hückel electrostatics for all pairs of charged residues. More details about this model can be found in Ref. [28]. For simplicity, other terms in the potential energy function due to angular or dihedral constraints are not included here.

Langevin dynamics simulations of the coarse-grained model are performed using LAMMPS-17Nov2016 [43] in an NVT ensemble at 300 K. These simulations are coducted in the low-friction limit with a small damping time constant (1 ps) to speed up convergence. The dielectric constant is set to that of water, 80, and the Debye screening length is set equal to 10 Å, which corresponds to the screening length at physiological salt concentration of 100 mM. Based on the results of all-atom simulations, PRMs are kept rigid in the PPII conformation as highlighted in Fig. 1A.

To study the effects of profilin binding on FH1 size, simulations were initialized by alignment of mouse profilin IIa to the FH1 PRMs using its bound crystal structure to mDia1-FH1(2PRM) as a reference (pdb accession code 2V8F) [32]. Simulations with profilin-actin complex bound to the PRM were initialized in a similar manner, with an additional step of cow beta-actin to mouse profilin IIa alignnment using the bound crystal structure to bovine profiling I as a reference (pdb accession code 2BTF) [47]. Individual profilin(-actin) units were kept rigid as in the original KH model [28]. Harmonic bonds were added to keep profilins bound to the FH1 PRMs and were treated exactly the same way as other bonds. These bonded pairs of residues are the ones for which H-bonds were identified in the crystal structure of profilin-mDia1 [32]. In the FH1 simulations with bound profilin(-actin), for greater than two PRMs occupied, the masses of the rigid profilin(-actin) domains was reduced (by a factor equal to the number of residues) to enhance the conformational sampling.

Radius of gyration and end to end distance are calculated from the backbone atoms for the AA simulations and from all beads for the CG simulations, which then allows the subsequent extraction of the asphericity.

The formin-bound barbed end simulations were initialized by alignment of the FH1 domains onto the FH2-bound barbed end model of [3], such that there were no gaps in sequence between the FH1 sequence used and the FH2. All actin subunits and the FH2-bound barbed end unit are made rigid. For Fig. 5, distances were defined as the distance between the center of masses (COM) of the PRM and the binding pocket of profilin with mDia1-FH1 as identifed in the crystal structure [32].

## 3 RESULTS

### FH1 Structure and Dynamics: All Atom Simulations

In order to study the structure and dynamics of FH1 domains, we performed all-atom simulations of a set of representative FH1 domains of formins which have been widely studied in prior experiments: mouse formins mDia1 (FH1 segment containing 6 out of total 14 PP regions as in [55], labeled mDia1-FH1(6PRM)) and mDia2, the two budding yeast formins Bni1 and Bnr1, and all three fission yeast formins Cdc12, For3, and Fus1 (Fig. 1). We used a protein force field (Amber99SBw) in combination with an optimized water model (TIP4P/2005) which was shown to be quite suitable for simulating unfolded proteins and IDPs [7, 26, 46]. Standard MD simulations provide dynamical information; however, the equilibration time of the longest FH1 domains studied here could be beyond the reach of MD simulations. We thus first compared the results of FH1 simulated with the enhanced sampling PTWTE method to MD simulations for two, mDia1 and Cdc12, out of the seven FH1 domains.

MD simulations required ~500 ns to give results in general agreement with PTWTE simulations (Fig. 2, S1). Neither mDia1-FH1(6PRM) nor Cdc12-FH1 develop any significant alpha helical or beta sheet structures in the PWTE and MD simulations, with the exception of a short segment in the middle of Cdc12-FH1 adopting a beta sheet turn with high propensity (Fig. S1). By contrast, the PRMs of both peptides have high propensity for PP conformation (Fig. 2), which is expected to be important for profilin binding. Indeed, the two PRMs in a crystal structure of profilin bound to a short mDia1-FH1 segment are primarily in the PPII conformation [32].

**FIGURE 2.**
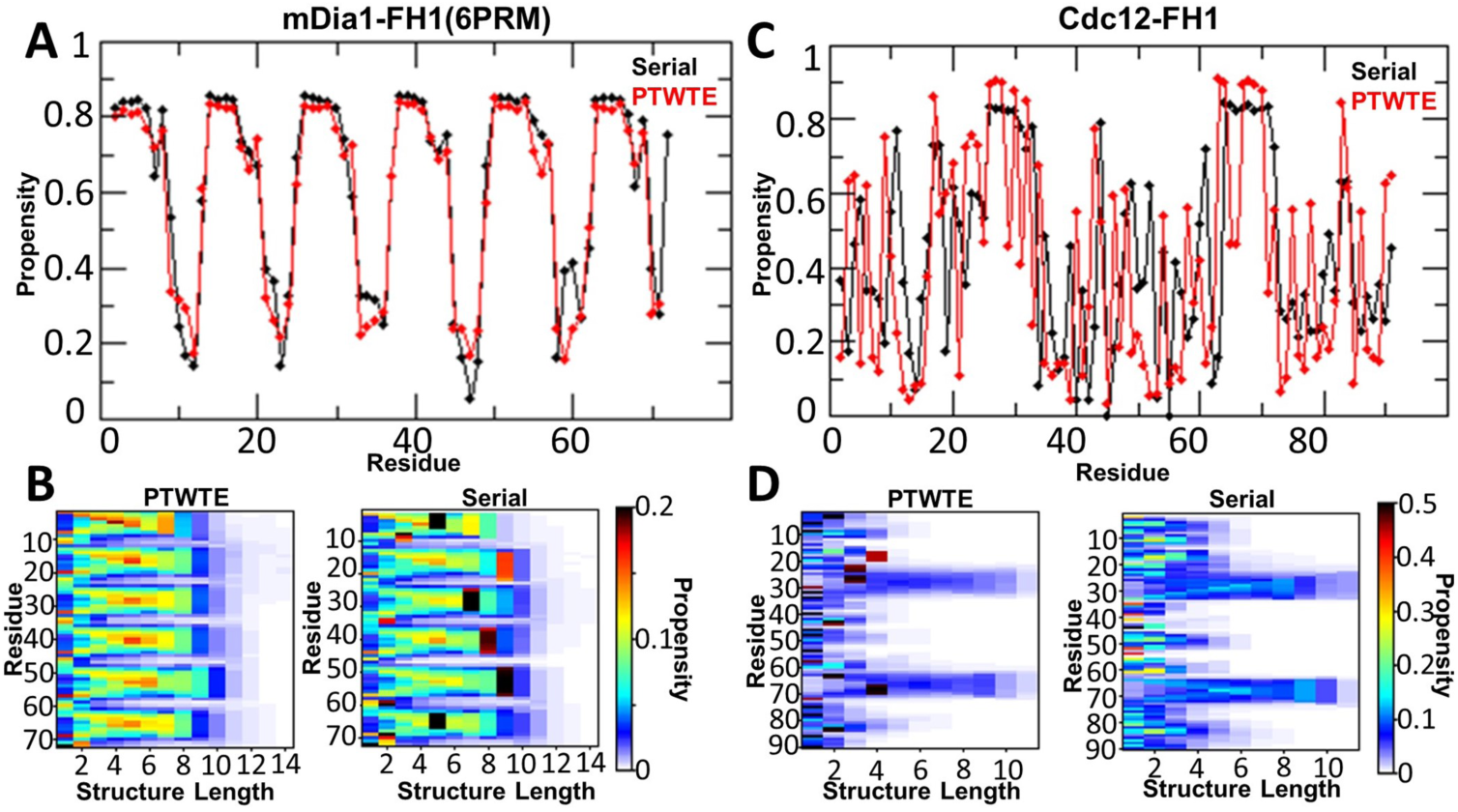
FH1 contains high propensity poly-L-proline helices, results from AA serial and PTWTE simulations. (A) and (C): Per residue propensity of PP of mDia1-FH1(6PRM) and Cdc12-FH1.(B) and (D): Poly-L-proline structure map, showing the propensity of a given residue to be in a given length PP structure for mDia1 and Cdc12, respectively.

This consistency motivated us to perform MD simulations for a similar timescale as for mDia1-FH1 and Cdc12-FH1 for the remaining FH1 domains. The PRMs of all FH1 domains contain high-propensity PP segments, notably even in regions with less than three successive prolines (Fig. S2). mDia2-FH1 has a PP stretch of variable length in its middle, with other regions of high PP propensity on either side: two closer to the N-terminus and one towards the C-terminus. Bni1-FH1 has four PP stretches, consistent with the number of profilin binding sites determined in prior experiments [13]. Four stretches are distinguished in Bnr1, up to five in For3 while Fus1-FH1 has more disorganized stretch close to its N-terminus. A small tendency for *α*-helix formation was observed near the C-terminus (adjacent to FH2) of Bnr1-FH1, Cdc12-FH1, and For3-FH1, and near the N-terminus of Bni1-FH1 (Fig. S3). No *β*-sheet structures were observed in these other FH1 domains (Fig. S4). Enhanced sampling techniques would be required to determine *α*-helix and *β*-sheet propensities with more confidence [15]; however, these were not performed here due to the computational cost.

The FH1 features described in the previous paragraph can be observed in supplementary Movies 1-7 of serial simulations. These show the higher rigidity of PP helices as well as heterogeneous structure/dynamics which is typical of unfolded proteins, with occasional transient alignment of interacting PP stretches and formation of short *α*-helical structures.

The size of the FH1 was quantified by measuring the distribution of the radius of gyration (*R*_*g*_), which had an average near 3 nm for mDia1 and Cdc12 in PTWTE and MD simulations (Fig. 3A,B). The average value of *R*_*g*_ for all FH1 MD simulations is shown in Fig. 3C, with distributions and time traces in Fig. S5. Prior analysis of experimental data on IDPs showed that, in addition to the number of residues, the most important parameters in determining the hydrodynamic radius *Rh* are net charge and proline content [34]. Using the values of Table 1, the “M&F-K” formula [34] gives *Rh* values comparable to *R*_*g*_ found in our simulations (Fig. 3), which highlights the importance of these parameters in the properties of FH1 domains as well. Given the complexities associated with the differences between *R*_*g*_ and *Rh* and the already good agreement between the two in our data, we do not attempt to interpret the differences, as there is a limited influence that is expected for most of the FH1 domains considered here [38]. We also note that the model of mDia1-FH1(6PRM) structure by Zhao *et al.* [55] predicted a similar *R*_*g*_ compared to our all-atom simulations (“ZLHL” in Fig. 3).

**FIGURE 3.**
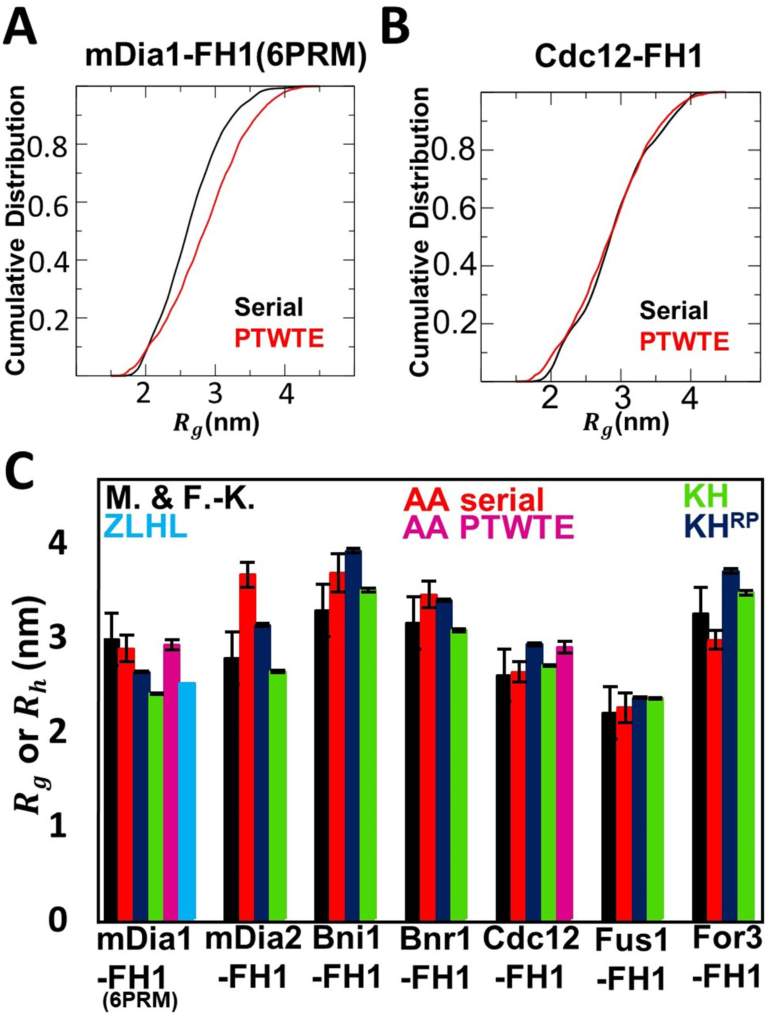
Size of FH1 is consistent with that of a typical IDP. (A) and (B): Cumulative distribution of radius of gyration of mDia1-FH1(6PRM) and Cdc12-FH1 in AA serial and PTWTE simulations. (C) Comparison of M&F-K empirical prediction of IDP hydrodynamic radius [34] to radius of gyration in AA serial, AA PTWTE, coarse-grained simulations using the KH and KH^RP^ models, and the ZLHL radius of gyration calculation for mDia1 [55].

**TABLE 1.**
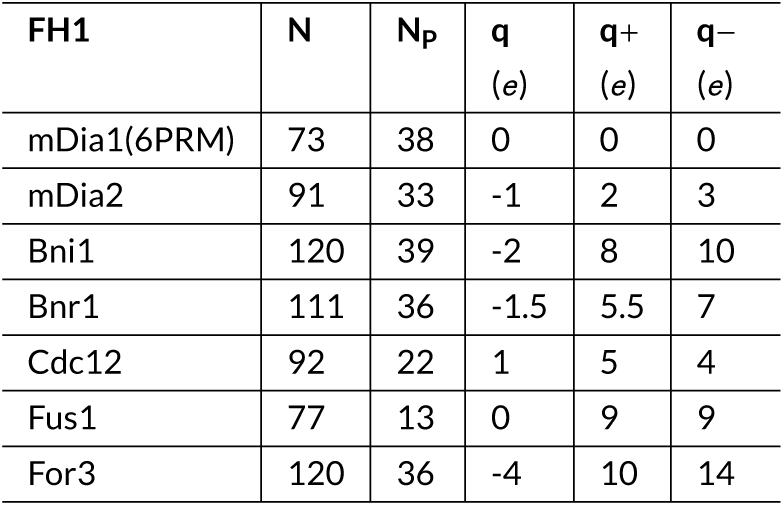
Relevant properties of FH1 domains studied in this work. Columns are: which FH1, number of residues (*N*), number of prolines (*N*_P_), net charge (*q*), positive (*q*+), and negative (*q*−) charges.

The relaxation time of *R*_*g*_ for the FH1 domains studied in this work was found to be in the order <50 ns (Fig. S5), comparable to the relaxation time of IDPs of similar length [37].

### FH1 Structure and Dynamics: Coarse-grained Model

All-atom simulations become computationally too costly when examining FH1 interactions with profilin and actin. Thus we used a coarse-grained model that accounts for the disordered FH1 configuration while maintaining sequence-specificity, using one bead per amino acid placed at the location of the C*α* atom and without explicit water. This bead retains the mass and charge of the amino acid, and can experience attractive or repulsive interactions with other beads based on the Miyazawa-Jerniagan pairwise interaction matrix. As a starting point, we used the KH model [28], see Materials And Methods. This model captures the sequence specificity but not the increased rigidity of PP helices. So we further made the PRMs of each FH1 (Fig. 1) explicitly rigid in a PPII configuration (model “KH^RP^”). Data were collected for at least 10 μs simulation time to ensure convergence (Movies 8-14). The KH^RP^ model captured the FH1 size measured in AA serial simulations (average deviation of 10.8%; Fig. 3). The rigidity of PP typically contributed to an increase of a few %, as observed by comparing the KH^RP^ to the KH model (average deviation from AA serial simulations of 12.1%; Fig. 3).

### FH1 Bound to Profilin and Profilin-Actin

We next examined how many profilins or profilin-actin complexes can bind to FH1, and how this binding expands FH1 size [11, 55]. FH1 could be highly occupied in cells where the concentrations of profilin and profilin-actin complex are tens of *μM* or higher, given that the binding affinity of profilin to mDia1-FH1(2PRM) is on the order of 10*μM*[32]. A detailed analysis would require a dynamic model of binding in which the ensemble-averaged occupancy of FH1 is determined by the bulk concentrations of profilin and actin. We took a simpler approach, considering that the dissociation time of profilin from long PP stretches is as long as 6 · 10_5_ ns (measured for Acanthamoeba profilin [2]) and profilin dissociation from actin occurs in seconds [52], both much longer than the estimated relaxation time of 50 ns for free FH1. Hence, using the coarse-grained KH^RP^ model, we treated profilin or profilin-actin as single rigid bodies permanently associated with specified PRMs and used mDia1-FH1(6PRM) as a reference FH1 (Fig. 4A). Specific occupancy was accomplished by adding bonds consistent with profilin binding to mDia1-FH1 as found by crystallography [32].

**FIGURE 4.**
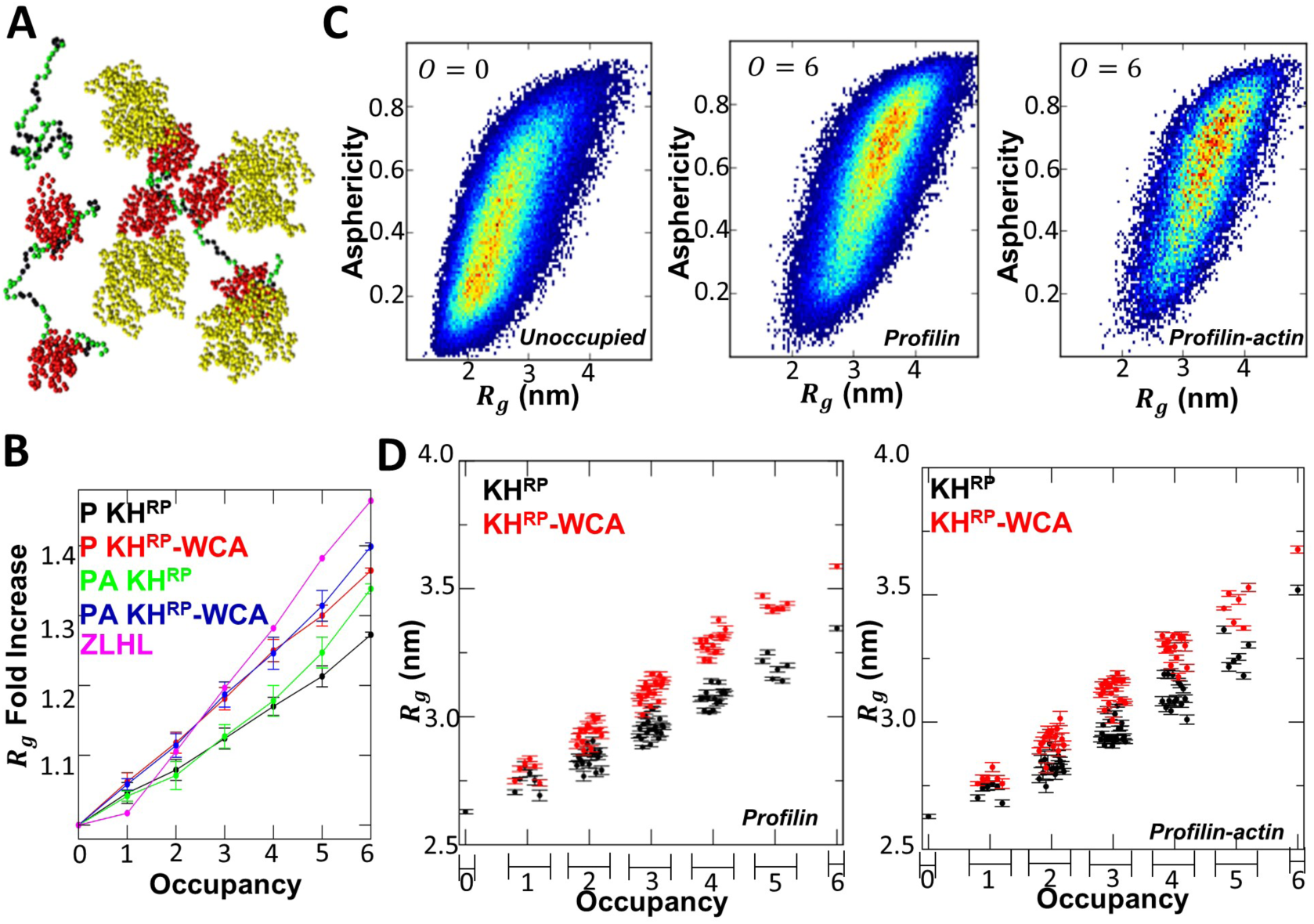
Weak expansion of FH1 upon profilin and profilin-actin occupancy. (A) Screenshots of typical conformations of unoccupied mDia1-FH1(6PRM) simulation and an example of a profilin-occupied and profilin-actin occupied mDia1 simulation. mDia1 PRMs colored in black, other mDia1 residues in green. Profilin colored in red and actin colored in yellow. For these snapshots the FH1 *R*_*g*_ and distance between each pair occupying profilin(-actin)s is within the FWHM of their respective distributions. (B) Fold increase in mDia1-FH1(6PRM) *R*_*g*_ from respective unoccupied *R*_*g*_ predictions for the KH^RP^ and KH^RP^-WCA models, as described in the main text, and comparison to ZLHL prediction [55]. Labels P and PA indicate profilin or profilin-actin occupancy. (C) KH^RP^ model predictions for *R*_*g*_ vs. asphericity distributions for unoccupied ( *O*= 1) and fully occupied ( *O*= 6) profilin or profilin-actin occupied mDia1-FH1(6PRM). (D) *R*_*g*_ predictions of the KH^RP^ and KH^RP^-WCA models for all possible profilin (left) or profilin-actin (right) mDia1-FH1(6PRM) occupancies. Horizontal ordering of data corresponds to the ordered set of occupancy configurations, binned according to total occupancy *O* =0 – 6.

**FIGURE 5.**
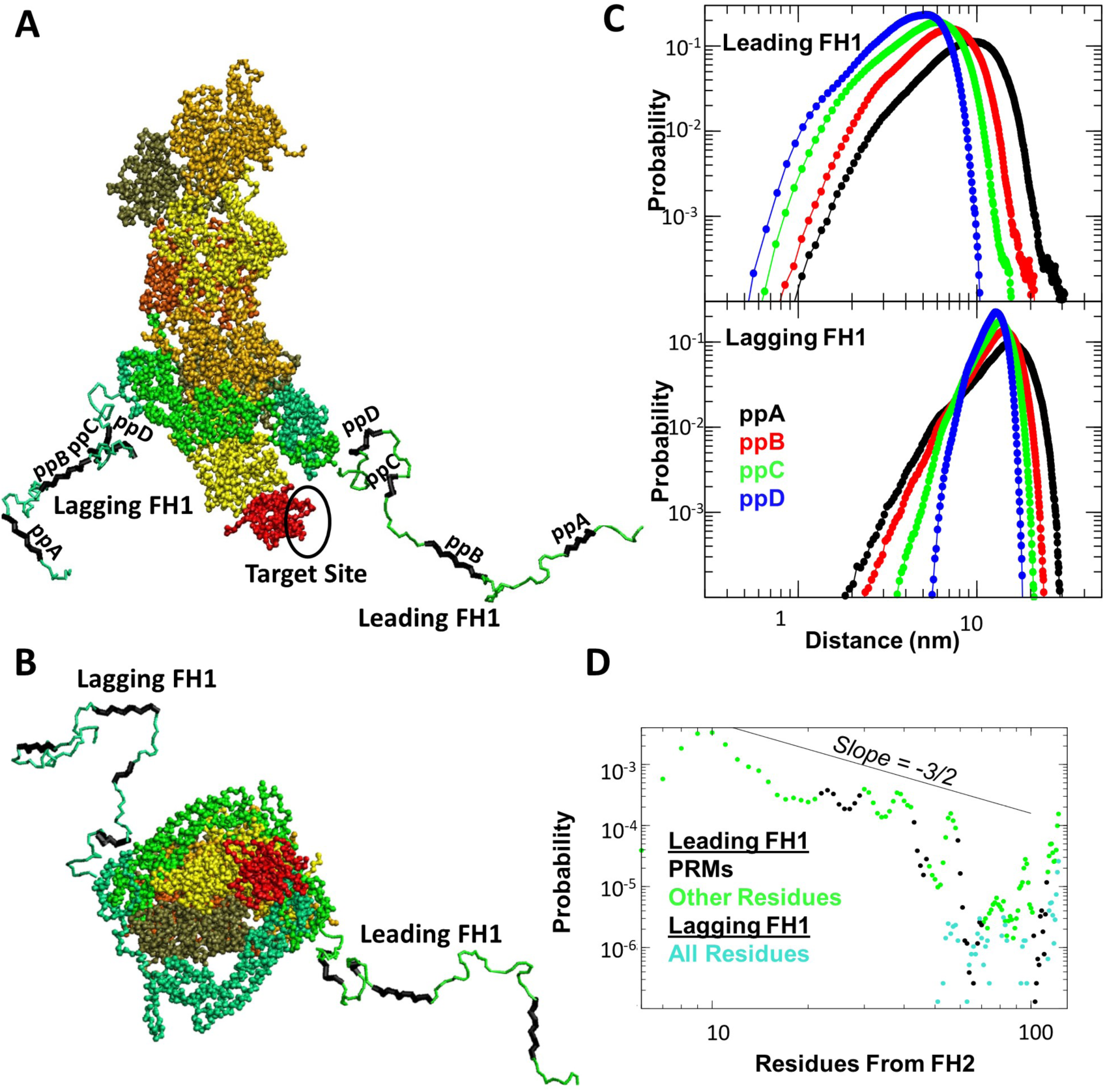
Coarse-grained simulations of FH1 as part of FH1-FH2 dimer associated with the barbed end of an actin filament with profilin bound to the terminal subunit. (A) Screenshot of typical conformation of Bni1-FH1-FH2 at the barbed end. FH1 and profilin colored as in Figure 4. The two FH2 domains and their associated FH1 segments colored in green and light blue, respectively. Actin subunits are colored in shades of yellow and orange. The shown FH1 conformations have *R*_*g*_ and COM distance between each FH1 PRM and the target binding site within the FWHM of their respective distributions (over the first 10 μs that are sufficiently long for equilibration). (B) Head-on view of the same frame as shown in (A). (C) COM distance distributions between each of PRMs for each FH1 and the target site. (D) FH1 per-residue probability to be within 1nm of the target site. Data points colored as in (A).

We ran simulations of all of the possible occupancy configurations of mDia1-FH1(6PRM) by profilin or profilin-actin complex (examples in Movies 15-16). This mDia1 sequence contains six PRMs which are approximately evenly distributed along the peptide chain. The ordered set of all occupancy configurations in terms of which sites (1 through 6, corresponding to the six PRMs) is: {{1}; {2}; {3}; …; {6}; {1, 2}; {1, 3}; {1, 4}; … {1, 6}; {2, 3}; {2, 4}; …;{5, 6}; {1, 2, 3}; {1, 2, 4}; …; {1, 3, 4}; … {1, 2, 3, 4, 5, 6}}, where each configuration has either profilin or profilin-actin occupying the sites. “Occupancy”, *O*, is defined as the number of bound objects there are on the FH1. There are six possible non-zero occupancies and 64 unique occupancy configurations. As a control case, we perform the same set of simulations, but using the purely repulsive (excluded volume interactions only) using Weeks-Chandler-Andersen (WCA) for the intermolecular interactions between profilin-FH1, actin-FH1, profilin-profilin, actin-actin, and actin-profilin (“KH^RP^-WCA” model). The latter gives a reference for the effect of intermolecular attractive or repulsive interactions. We show results using profilin II but similar results apply for the more highly charged profilin I, consistent with there being no difference between the two in mDia1-mediated actin polymerization [12]. To ensure sufficient sampling of bound molecule distributions, data were collected for at least 5μs simulation time for occupancy greater than 2, and at least 10μs simulation time for other simulations.

We found that profilin or profilin-actin complex can be added to all six sites of mDia1-FH1(6PRM). The change of FH1 shape due to binding was quantified using the *R*_*g*_,the end-to-end distance *R*_*ee*_, and the asphericity of FH1 (Fig. 4 and S6). Consistent with the expectation, expansion of FH1 is seen upon increase of profilin occupancy (Fig. 4B-D, S6). FH1 expansion depends slightly more strongly on profilin-actin occupancy than on profilin occupancy (Fig. 4B-E, S6). For each individual occupancy configuration, the KH^RP^-WCA model experiences greater expansion than KH^RP^. The asphericity and *R*_*g*_ are correlated, showing that FH1 becomes elongated as it expands through binding (Fig. 4C). The increase in FH1 size depends slightly on where profilin or profilin-actin is bound along the FH1. For occupancies *O* < 3, configurations having occupied sites closer to the middle of FH1 are more expanded compared to configurations with the terminal sites occupied (for *O* =1 this can be seen in Fig. 4, where the data points are ordered according to the occupied site). This is most likely due to the larger number of excluded steric contacts when the middle of FH1 is occupied as compared to the terminal region.

The dependence of *R*_*g*_ on profilin occupancy for our models is slightly weaker than previously predicted by ZLHL [55] (Fig. 4). In this prior prediction, the equilibrium ensemble was determined by simple counting of geometric constraints, i.e., whether a specific conformation for a given occupancy configuration contained any clashes. If no clashes were found, the conformation was considered part of the equilibrium ensemble. This procedure may have excluded states which have attractive contacts between atoms, such as sidechains from two different residues. Another possible origin of the disccrepancy with ZLHL is poor sampling of extended FH1 configurations since the occupied FH1 ensemble was taken by excluding configurations from the unoccupied ensemble. Bryant *et al*. also predict expansion of FH1 upon multiple profilin-actin occupancy by excluding steric interactions in a uniform freely-jointed chain model; in simulations with mDia1-FH1 containing 14PRMs, we find a similar expansion of 20% for the same 3 occupied sites [11], using KH^RP^.

### Transfer of Profilin-Actin by FH1 to the Barbed End

Finally, we considered FH1 in the context of its function to transfer profilin-actin to the barbed end. A recent work utilized available crystal structures and actin filament models to build an all-atom model of the actin filament with a dimer of the Bni1 FH2 domain bound at the barbed end [3]. We used this model of FH2-bound Bni1 to test whether a newly added actin subunit can be transfered as profilin-actin to the barbed end through one of the FH1 domains of a Bni1 FH1-FH2 dimer. This newly-added actin subunit bound to profilin (aligned to actin using the profilin-actin crystal structure [47]) is shown in Fig. 5A,B. While the precise mechanism of FH2 stepping is not fully resolved, the configuration in Fig. 5 is, most likely, the configuration after addition of a new profilin-actin subunit and prior to the stepping forward of the lagging FH2 domain of the dimer, in preparation to accept the next subunit [41]. Specifically, this corresponds to the addition of a new subunit in the “stair-stepping model” [31].

As with the occupancy simulations of Fig. 4, studying the kinetics of the profilin-actin transfer mechanism explicitly would require a detailed model of binding and polymerization [39, 48]. We take here a simpler approach by asking if either FH1 domain can reach out and with what probability to the polyproline binding pocket of profilin at the terminal actin subunit. Calculating the equilibrium contact probabilty of each FH1 PRM to the target profilin site would provide us with an estimate of the profilinactin transfer for the correspondig PRM. This reasoning assumes that the FH1 profilin-actin transfer rate is proportional to the equilibrium probability of finding FH1-bound profilin-actin at the barbed end [52].

To construct the Bni1 FH1-FH2 dimer at the barbed end, we coarse-grained the AA model of Baker *et al.* [3] to one bead per residue and appended FH1 domains to each respective FH2 after adding two missing residues between the sequence for Bni1 FH1 of Fig. 1 and the Bni1 FH2 sequence in the structure of [3]. Actin, profilin and FH2 were treated as rigid objects while FH1 was allowed to move freely, interacting with each residue using the KH model.

Performing MD simulations for 80-100μs total simulation time (Movie 17-18), we calculated the distance between the binding pocket of profilin and each of the four presumed Bni1 profilin binding sites labeled ppA-ppD [41]. These sites correspond to the four regions of high PP propensity of Fig. S2. Since ppD (r93-101) has somewhat lower PP propensity, we left it flexible while the other PRMs were kept in the PPII configuration. Note that we assume the binding pocket on profilin is the same for the FH1 of Bni1 as for mDia1 in Fig. 4.

We find that only the FH1 labeled “leading” is in position to approach the profilin site and make frequent transfer attempts while the “lagging” FH1 is significantly more distant (Fig. 5C). The probability of finding a leading FH1 residue in close proximity to the profilin pocket decreases with site distance from the FH2 domain as suggested in [52], however notably this trend is reversed for the lagging FH1 (see probabilities for distances < 1nm in Fig. 5C). A minimum of 5-6 residues away from the FH2 are needed for leading FH1 to reach the profilin binding site, with the 10*^th^* residue having the highest probability (Fig. 5D). For the more distant residues, the equilibrium contact probability decays approximately as *n*^−3/2^, ideal random walk [52]), however the decay is not monotonic as a result of interactions and/or PP rigidity (Fig. 5D).

Removing the terminal profilin-actin of Fig. 5 and placing profilin on the terminal actin of the other protofilament leads to a configuration with the lagging FH1 being closer to the PRM profilin binding site (Fig. S7). FH1 binding to that site might occur during the formin cycle though this is less likely to lead to polymerization or depolymerization of this more hidden profilin-actin terminal subunit.

An estimate of the rate with which PRMs reach to the profilin-binding site can be obtained by assuming that the rate is proportional to the equilibrium contact probability [52]. Over a simulation time of 76μs, the number of closure events for site ppD is on the order of 1000. Taking into account that the relaxation time for *R*_*g*_ in the coarse-grained simulations is shorter by a factor 2-3 than in the AA simulations (because of the small drag coefficient used to speed up thermodynamic convergence) yields a contact rate of the order of 10^5^*s*_-1_. This value is consistent with the estimate of 10^4^*s*^−1^ for profilin-actin delivery from the closest FH1 profilin binding site in the direct transfer model [52]

## 4 DISCUSSION AND OUTLOOK

Application of an all-atom force field which well-represents both the size and residual secondary structure of disordered proteins suggested the general picture of FH1 as a segment with flexible domains intermingled with regions of more rigid high-propensity poly-L-proline helices in the proline-rich regions. The size of several FH1 domains we considered, measured by the radius of gyration *R*_*g*_, was consistent with the empirical M&F-K formula for IDP hydrodynamic radius *R_h_* [34]. The high rigidity of the polyproline helix, as compared to the persistence length of a typical IDP (a few residues [25]) and the “blockiness” of the prolines has a small but noticeable effect on FH1 size. Results from our coarse-grained model (with and without rigid PRMs) showed that this increase is typically a few per cent for the FH1s considered in this study.

Zhao *et al.* proposed that multiple profilin-actin binding to FH1 leads to a “cooperative jack model of random coil-to-elongation transition” [55]. With regard to the simultaneous binding of profilin or profilin-actin molecules to multiple PRMs (e.g., mDia1), we show that there are no steric constraints excluding the possibility of multiple occupancy. The development of an F-actin-like arrangement of multiple bound profilin-actin, which is presumably needed for this proposed cooperative transition, was not apparent in our simulations (Movie 16). As interactions between actin molecules are quite weak in the KH model used here, one should first reparameterize the model using experimental data to capture the expected binding affinity as well as the native complex structure [29]. We intend to do this in future work.

Prior work hypothesized that FH1 domain favors delivery of actin to the nearest long-pitch helix [13]. To address this, we simulated the Bni1-dimer associated barbed end and find that indeed “leading” FH1 strand of the Bni1-dimer is much more likely to reach the profilin-binding site (to transfer profilin-actin) than the “lagging” strand. The estimated FH1 closure rate further supported the direct transfer mechanism [17, 18, 52].

Most importantly, the coarse-grained model presented here can provide accurate information on FH1 (with input from all-atom simulations) size which is highly sequence specific in contrast to the existing computational models. Therefore, it should serve as a useful tool to study formin-mediated actin polymerization because of its unique ability to distinguish the various roles and effects of different formins from the amino-acid level that existing models are unable to do.

## 5 ACKNOWLEDGEMENTS

We thank Professor Greg Voth for providing the initial structures of Bni1-FH2 bound to actin. BH thanks Greg Dignon for useful discussions on simulation methodology and coarse-grained models. BH and DV were supported by National Institutes of Health Grant R01GM114201. GHZ and JM were supported in part by National Institutes of Health Grant R01GM118530. This research used resources of the National Energy Research Scientific Computing Center, a DOE Office of Science User Facility supported under Contract No. DE-AC02-05CH11231. Use of the high-performance computing capabilities of the Extreme Science and Engineering Discovery Environment (XSEDE), which is supported by the National Science Foundation, project no. TG-MCB120014 and TG-MCB160144, is also gratefully acknowledged.

## Movie Captions

Movie 1: AA simulation of mDia1-FH1(6PRM). 30fps. Each frame 600ps simulation time.

Movie 2: AA simulation of mDia2-FH1. 30fps. Each frame 600ps simulation time.

Movie 3: AA simulation of Bni1-FH1. 30fps. Each frame 600ps simulation time.

Movie 4: AA simulation of Bnr1-FH1. 30fps. Each frame 600ps simulation time.

Movie 5: AA simulation of Cdc12-FH1. 30fps. Each frame 600ps simulation time.

Movie 6: AA simulation of Fus1-FH1. 30fps. Each frame 600ps simulation time.

Movie 7: AA simulation of For3-FH1. 30fps. Each frame 600ps simulation time.

Movie 8: CG simulation of mDia1-FH1(6PRM). 30fps. Each frame 5.5ns simulation time.

Movie 9: CG simulation of mDia2-FH1. 30fps. Each frame 5.5ns simulation time.

Movie 10: CG simulation of Bni1-FH1. 30fps. Each frame 5.5ns simulation time.

Movie 11: CG simulation of Bnr1-FH1. 30fps. Each frame 5.5ns simulation time.

Movie 12: CG simulation of Cdc12-FH1. 30fps. Each frame 5.5ns simulation time.

Movie 13: CG simulation of Fus1-FH1. 30fps. Each frame 5.5ns simulation time.

Movie 14: CG simulation of For3-FH1. 30fps. Each frame 5.5ns simulation time.

Movie 15: CG simulation of mDia1-FH1(6PRM) with profilin occupying two PRMs. 7fps. Each frame 11.5ns simulation time.

Movie 16: CG simulation of mDia1-FH1(6PRM) with profilin-actin occupying four PRMs. 7fps. Each frame 11.5ns simulation time.

Movie 17: CG simulation of Bni1-FH1-FH2 bound barbed end. 30 fps. Each frame 5.5ns. For convenience, barbed end is kept still in the movie.

Movie 18: Same as Movie 17, but a top view.

---

**FIGURE S1.**
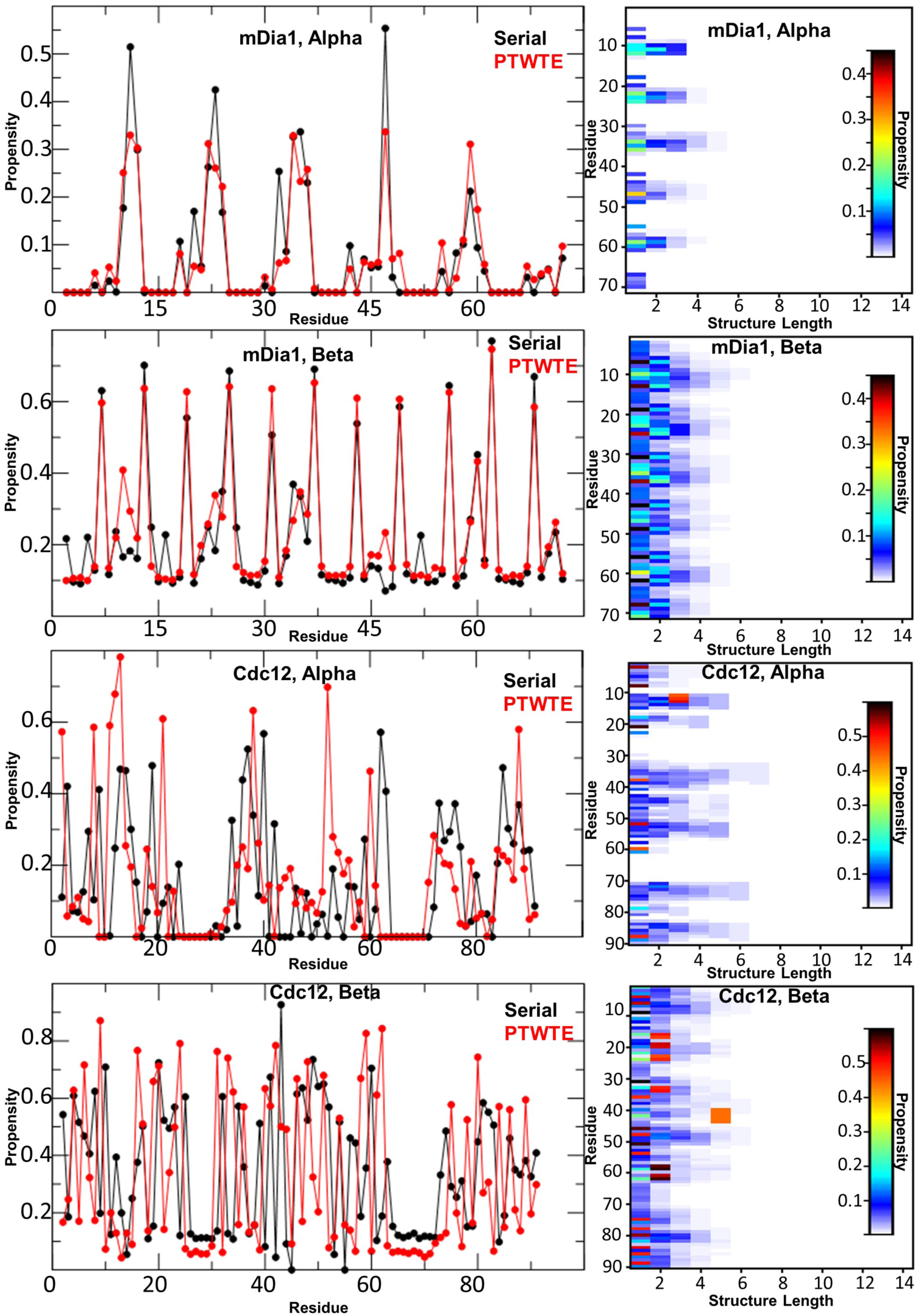
All-atom PTWTE simulations reveal little to no alpha or beta type structures in the **FH1** domains shown in Fig.1A of the main text. Secondary structure is classified by backbone dihedral angles.

**FIGURE S2.**
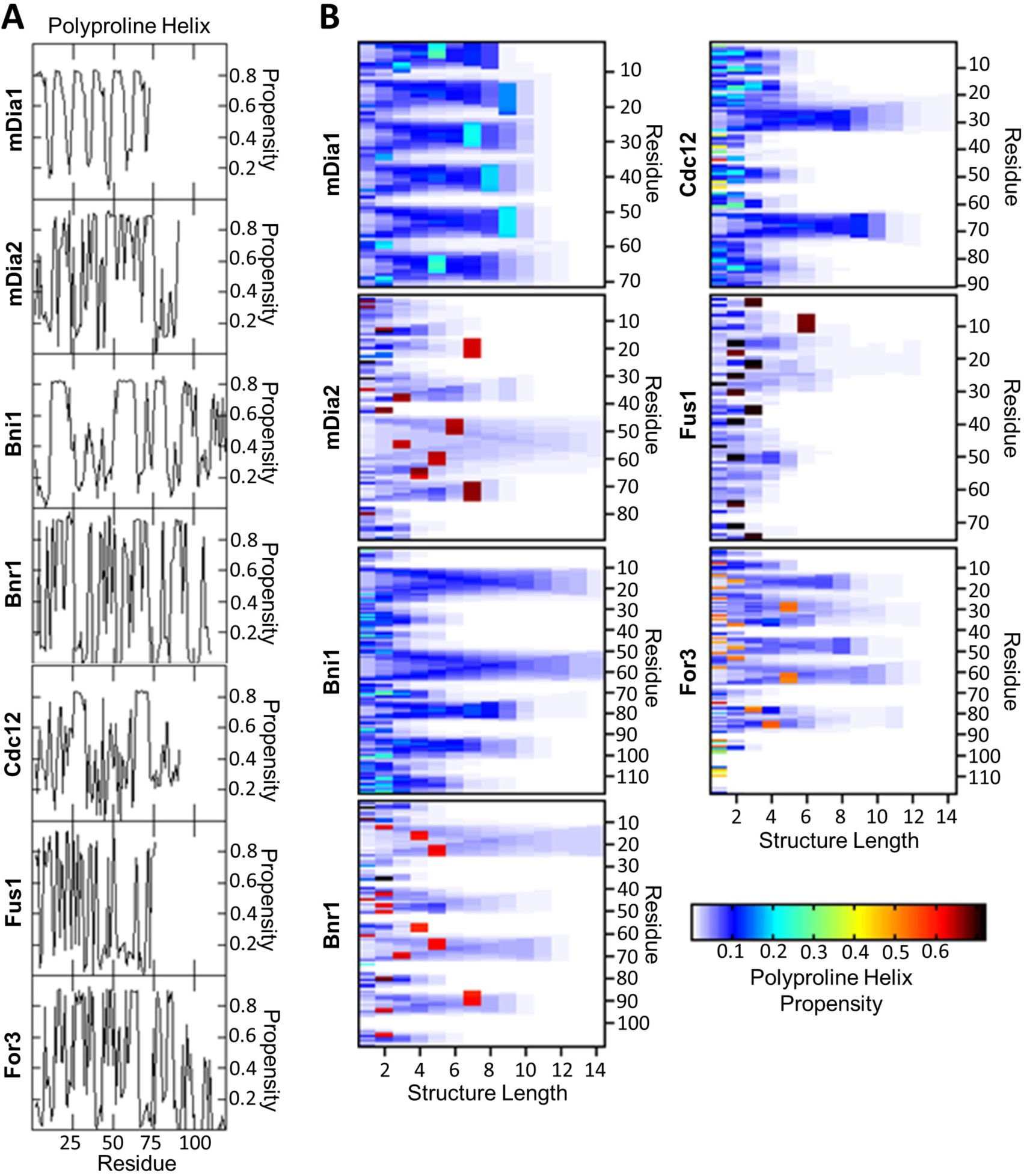
All-atom serial simulations reveal PP helices in the FH1 domains of Fig. 1A of the main text. **A**:Total per-residue propensity. **B**: Per-residue and structure length propensity for polyproline helix. Secondary structure is classified by backbone dihedral angles.

**FIGURE S3.**
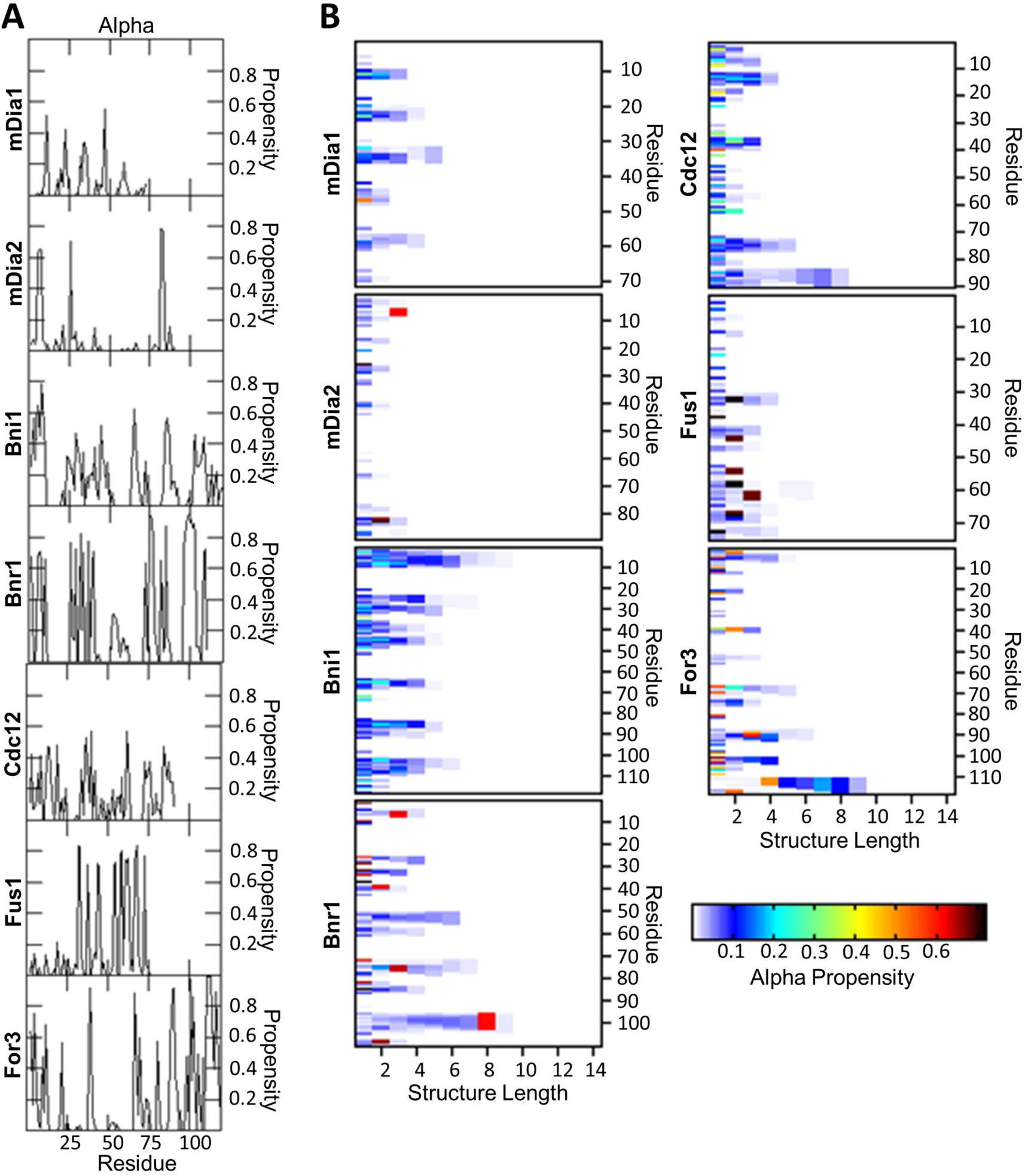
All-atom serial simulations reveal little to no alpha helical structures in the FH1 domain. **A**: Total per-residue propensity. **B**: Per-residue and structure length propensity for alpha helices. Secondary structure is classified by backbone dihedral angles.

**FIGURE S4.**
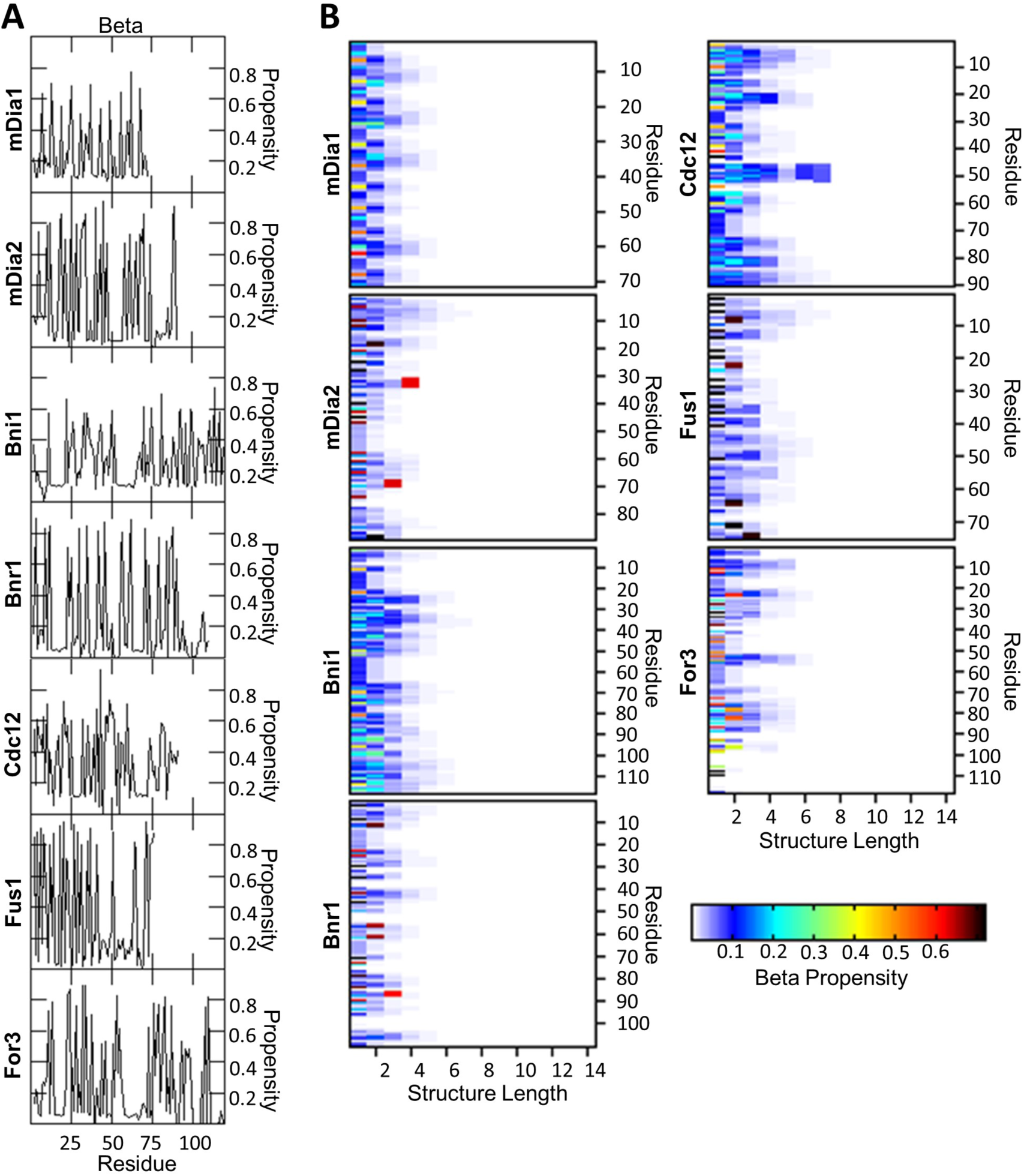
All-atom serial simulations reveal little to no beta type structures in the FH1 domain. **A**: Total per-residue propensity. **B**: Per-residue and structure length propensity for beta structures. Secondary structure is classified by backbone dihedral angles.

**FIGURE S5.**
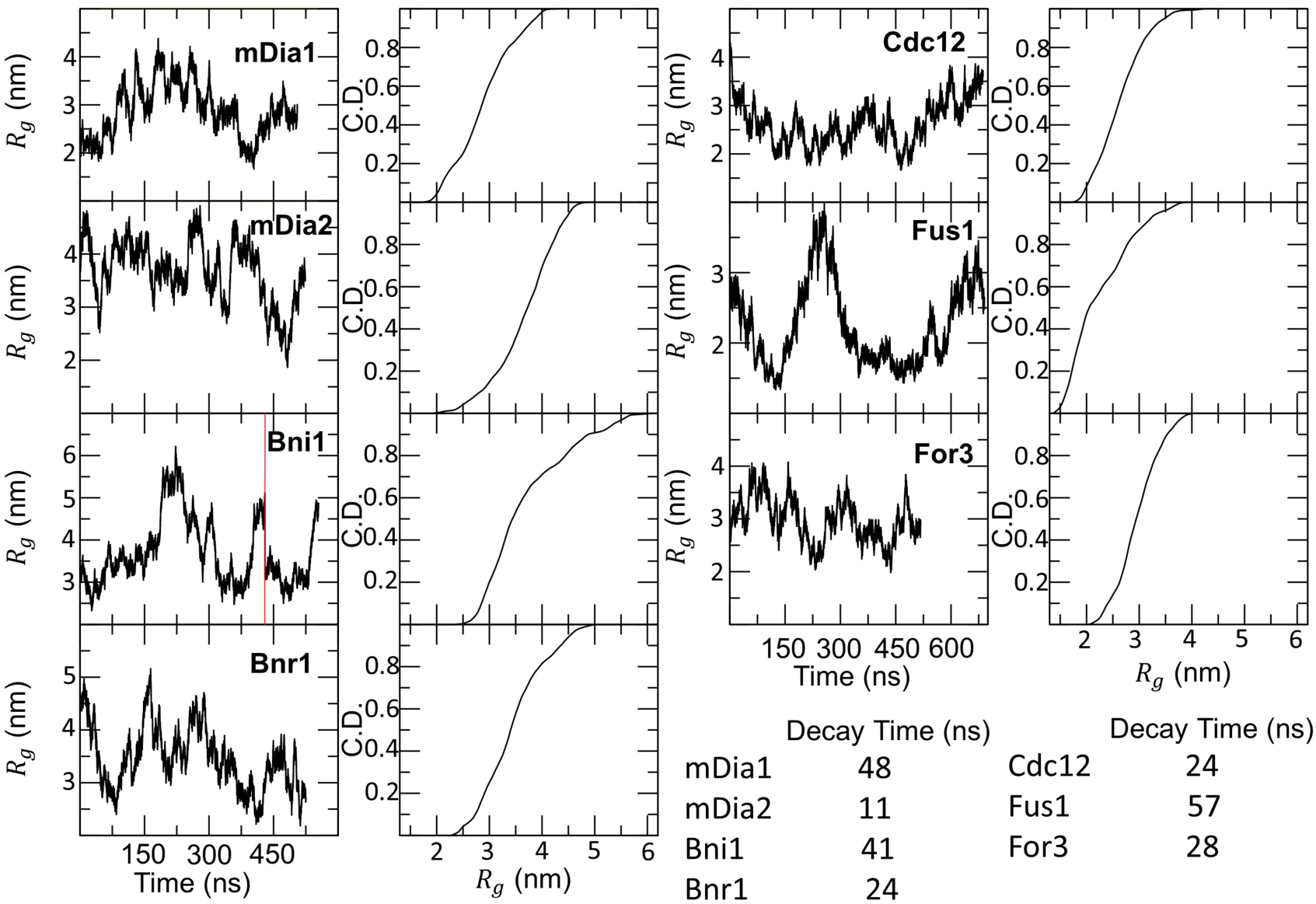
*R*_*g*_ time traces and distributions for the all-atom serial simulations of the FH1 domains showin in Fig. 1A of the main text. The red line in Bni1 represents where the simulation was cut off and resolvated in a larger box because continuation without this resulted in significant periodic image interaction. Decay times are the first time that the autocorrelation function of *R*_*g*_ drops below 1/ *e.* Decay time for Bni1 calculated for simulation before resolvated in larger box.

**FIGURE S6.**
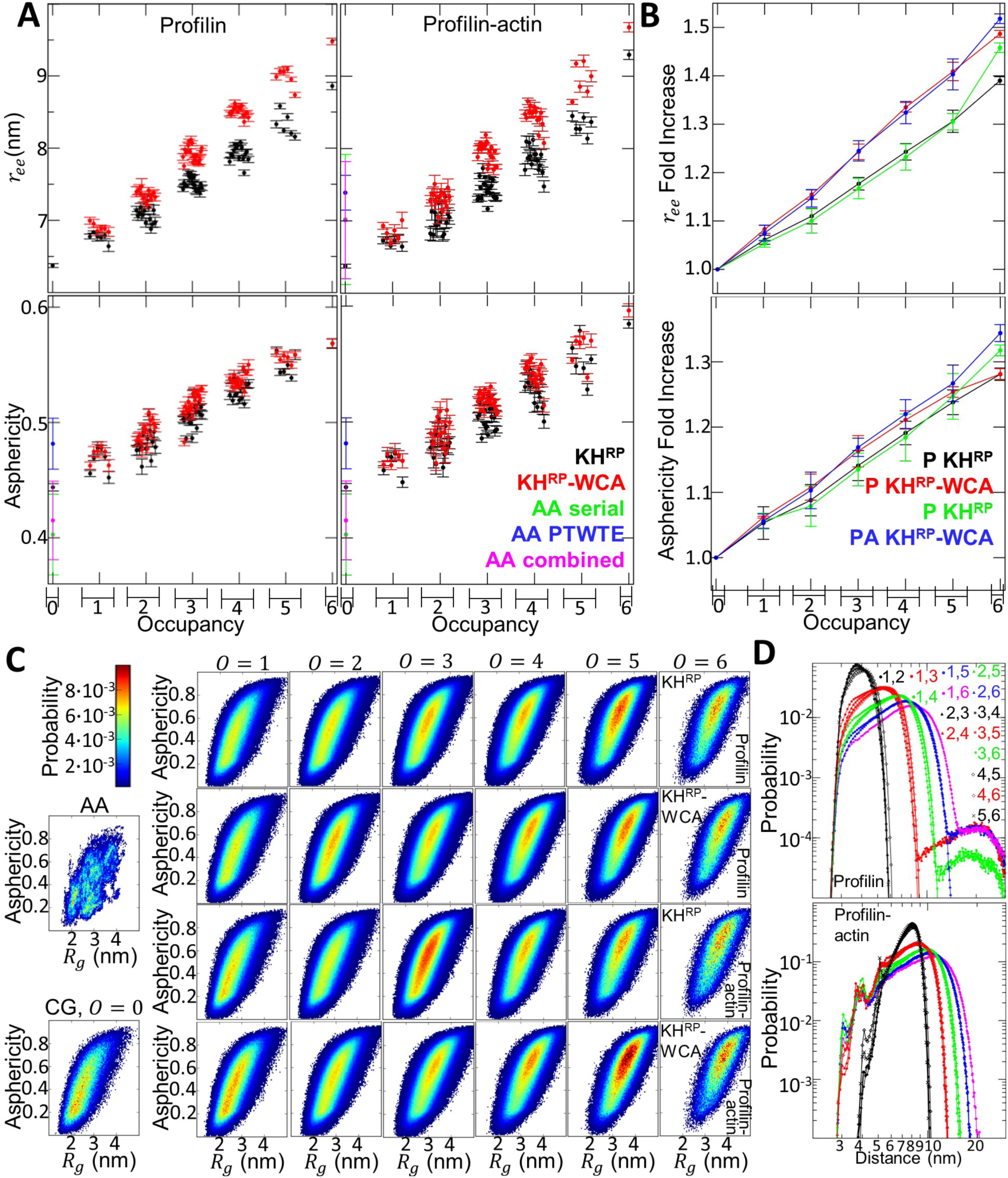
Additional Details on Occupied FH1 Size. **A**: Average end to end distance and asphericity, plotted in the same way as for *R*_*g*_ in Figure 5D. **B**: End to end distance and asphericity fold increase, plotted in the same way as for *R*_*g*_ in Figure 5B. **C**: Additional *R*_*g*_ vs. asphercity maps. Left column: unoccupied combined all-atom and unoccupied coarse-grained maps. The 4×6 block shows distributions over all simulations for a given occupancy for a given model for occupying profilin or profilin-actin. **D**: Distance distributions between occupying profilin or profilin-actin. Distributions are over all simulations containing a given pair of occupying profilin or profilin-actin.

**FIGURE S7.**
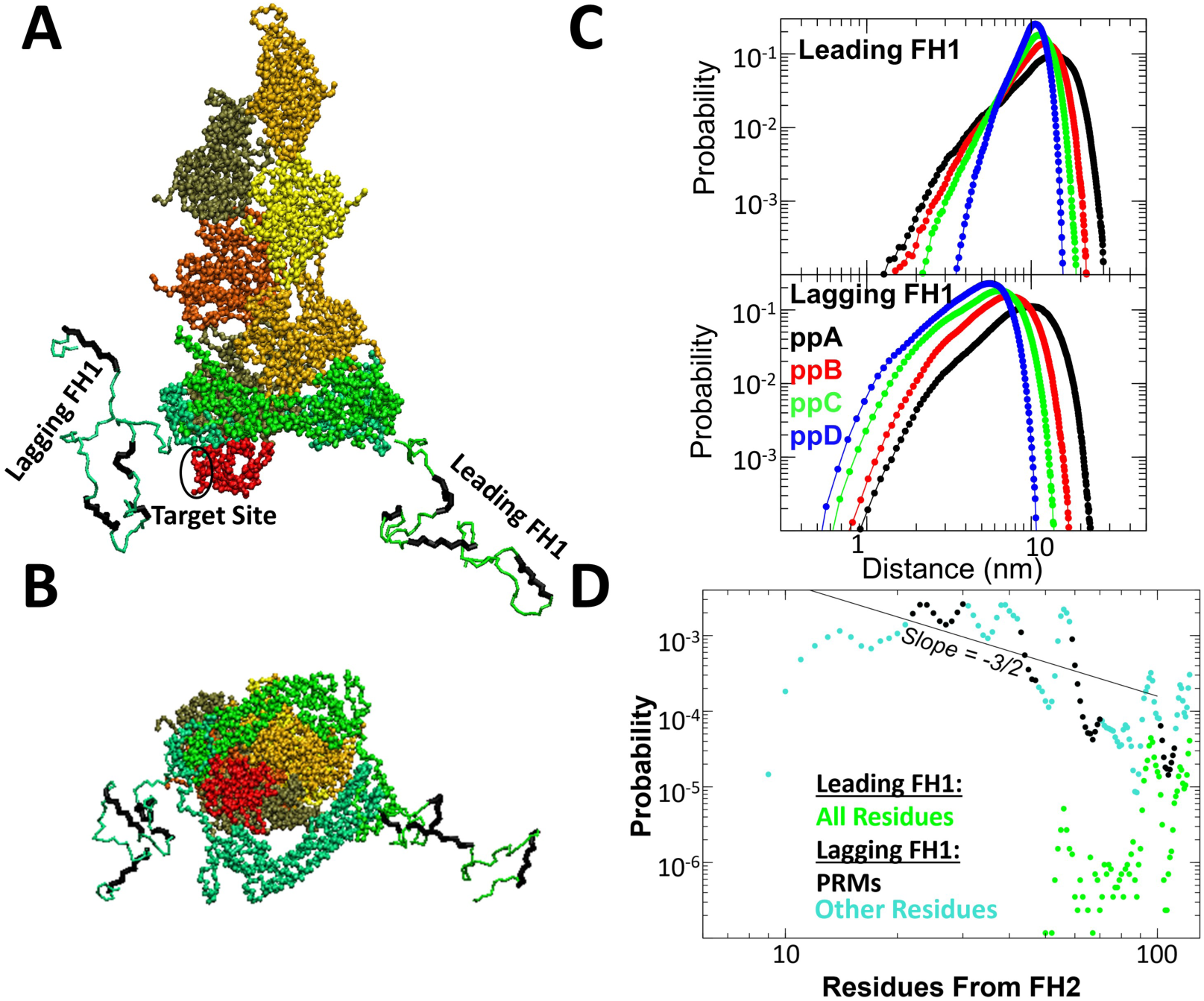
Coarse-grained simulations of FH1 as part of Bni1 FH1-FH2 dimer associated with the barbed end of an actin filament, with one less actin subunit at the barbed end compared to Fig. 5 of the main text. **A,B:** Screenshot of typical simulation configuration. This simulation is the same as that of Fig. 5, except that the most-terminal actin subunit has been removed, and profilin placed on the next actin subunit in the filament (i.e. the last subunit on the other protofilament). **C:** Distance distributions of FH1 PRMs to the target site on profilin (same as in Fig. 5) show the lagging FH1 is now favored for closure. **D:** Per-residue probability of closure for each FH1.

